# SARS-CoV-2 desensitizes host cells to interferon through inhibition of the JAK-STAT pathway

**DOI:** 10.1101/2020.10.27.358259

**Authors:** Da-Yuan Chen, Nazimuddin Khan, Brianna J. Close, Raghuveera K. Goel, Benjamin Blum, Alexander H. Tavares, Devin Kenney, Hasahn L. Conway, Jourdan K. Ewoldt, Sebastian Kapell, Vipul C. Chitalia, Nicholas A. Crossland, Christopher S. Chen, Darrell N. Kotton, Susan C. Baker, John H. Connor, Florian Douam, Andrew Emili, Mohsan Saeed

## Abstract

SARS-CoV-2 can infect multiple organs, including lung, intestine, kidney, heart, liver, and brain. The molecular details of how the virus navigates through diverse cellular environments and establishes replication are poorly defined. Here, we performed global proteomic analysis of the virus-host interface in a newly established panel of phenotypically diverse, SARS-CoV-2-infectable human cell lines representing different body organs. This revealed universal inhibition of interferon signaling across cell types following SARS-CoV-2 infection. We performed systematic analyses of the JAK-STAT pathway in a broad range of cellular systems, including immortalized cell lines and primary-like cardiomyocytes, and found that several pathway components were targeted by SARS-CoV-2 leading to cellular desensitization to interferon. These findings indicate that the suppression of interferon signaling is a mechanism widely used by SARS-CoV-2 in diverse tissues to evade antiviral innate immunity, and that targeting the viral mediators of immune evasion may help block virus replication in patients with COVID-19.

---

SARS-CoV-2, the virus behind the ongoing COVID-19 pandemic, has claimed more than one million human lives in a short time span of less than a year^1^ (https://coronavirus.jhu.edu/map.html). The virus primarily infects lungs, causing acute respiratory distress syndrome (ARDS) and respiratory failure^2^, which are one of the leading causes of death in COVID-19 patients. However, as more tissue specimens from infected and/or deceased individuals have become available and are probed for the presence of viral proteins or particles, there is increasing appreciation that SARS-CoV-2 can target multiple organs. Evidence for virus replication has so far been found in the intestine^3,4^, liver^5,6^, kidney^7,8^, heart^9,10^, and brain^11,12^.

The molecular mechanisms that govern the ability of SARS-CoV-2 to manipulate diverse cellular environments and navigate various body organs are unknown. One way to identify these mechanisms is to conduct a comprehensive survey of cellular pathways disrupted by the virus in distinct cell backgrounds. This however would require human-derived cell models that allow efficient and synchronized viral infection so as to generate a high-confidence catalog of virus-induced changes. Such systems are currently lacking, and as a result, a large number of SARS-CoV-2 studies, including a recent genome-wide CRISPR screen to identify the viral essentiality factors^13^, have been carried out in non-human cells.

To catalog cellular pathways broadly targeted by SARS-CoV-2, we generated a panel of 17 phenotypically diverse human cell lines that represented various body organs and supported high levels of SARS-CoV-2 infection. We leveraged this unique panel of cell culture models to profile proteomic responses to infection in cells originating from lung, liver, intestine, kidney, heart, and brain. This led to identification of cellular proteins and pathways widely targeted by the virus across cell types. Notable among these pathways was the JAK-STAT signaling cascade, the key component in the interferon response pathway. We performed extensive validation and functional studies, both in cell lines and pluripotent stem cell-derived cardiomyocytes, to show that SARS-CoV-2 rapidly disables the JAK-STAT pathway and consequently desensitizes host cells to interferon treatment. In line with this, the chemical inhibition of JAK-STAT signaling enhanced SARS-CoV-2 infection. These results uncover an immune evasion strategy of SARS-CoV-2 to create a favorable environment for its replication in diverse tissue types and highlight virus-mediated immune antagonism as a target for therapeutic intervention.

## RESULTS

### A large number of human cell lines are resistant to SARS-CoV-2 infection

Seventeen human cell lines derived from lung, intestine, heart, kidney, liver, brain, and vasculature (**Fig. 1a**) were infected with SARS-CoV-2 at a low multiplicity of infection (MOI) of 0.1 to allow multiple cycles of virus replication and spread. Immunofluorescence (IF) analysis showed that nine of the 17 cell lines were completely resistant to infection (**Fig. 1b**). Of the remaining eight, three, Calu-3, HepG2, and Caco-2 cells reliably supported measurable virus infection at 24 hours post-infection (hpi), although at levels much lower than those observed for the widely used Vero E6 cells (>10%, compared to 73% for Vero cells). The number of virus-infected cells further increased to around 35-42% for Calu-3 and Caco-2 cells at 72 hpi, and the infection also became detectable in HK-2 (48%), HuH-6 (47%), HuH-7 (22%) and Huh-7.5 (27%) cells at that time.

**Fig. 1:**
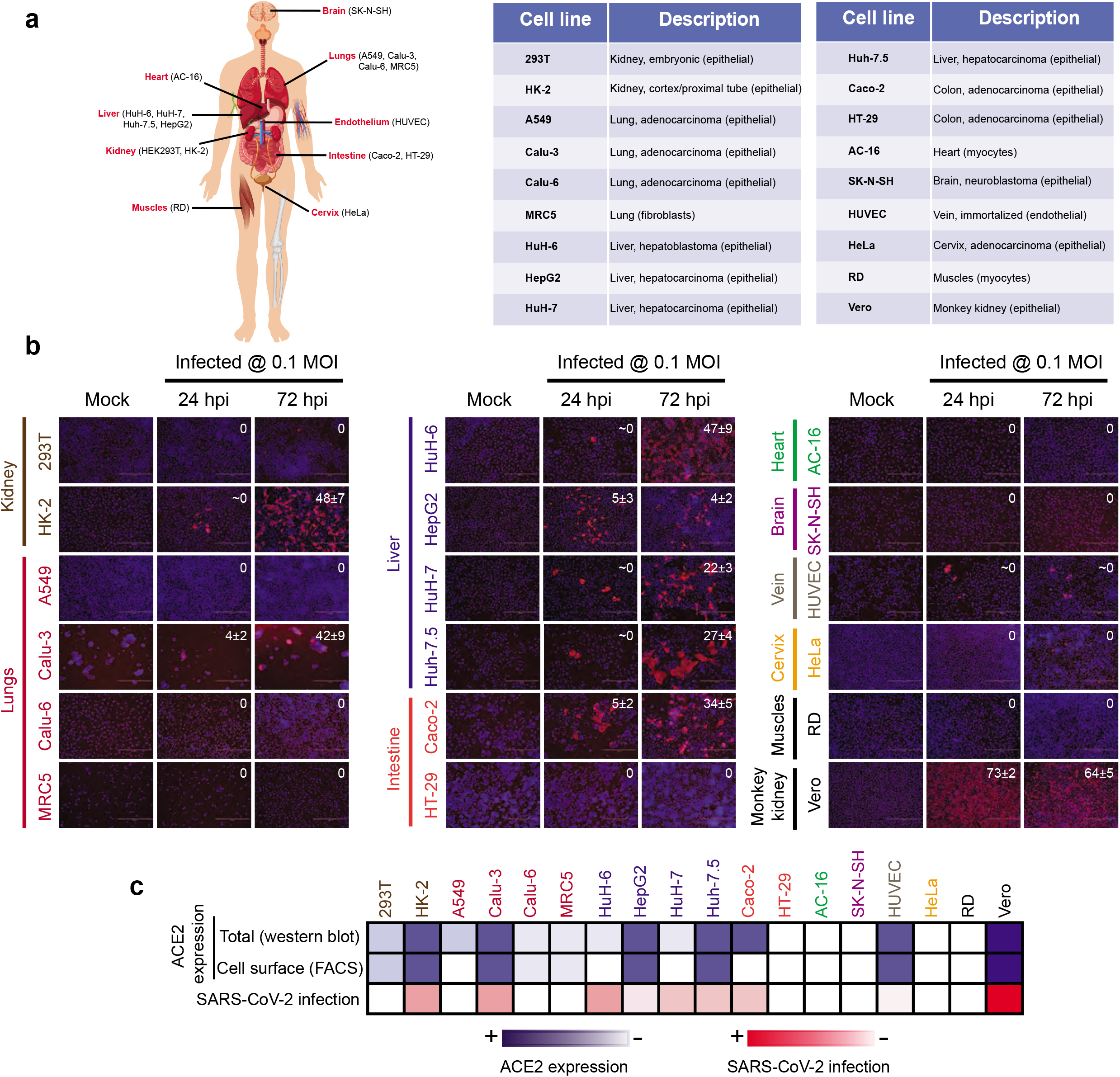
Most human cell lines supported no or minimal infection by SARS-CoV-2. **a**, The cell lines and their tissues of origin are shown. The detailed description of each cell line is provided in the table. **b**, The indicated cells were infected with SARS-CoV-2 at an MOI of 0.1 and stained with the viral nucleocapsid (N) protein (red) at 24 and 72 hpi. The nuclei were counterstained with DAPI. The mean percentage of positive cells ± standard deviation of three biological replicates is shown. The cell names are colored differently according to their tissues of origin (n = 3). **c**, Total abundance of ACE2 in cells and its cell surface-associated fraction was measured by Western blot and flow cytometry, respectively. The shades of blue indicate the intensity of the ACE2 signal, whereas the shades of red reflect the SARS-CoV-2 infection efficiency in each cell line. The data of Western blot and flow cytometry experiments are presented in **Extended Data Fig. 1**.

When we analyzed ACE2 expression through Western blot (**Extended Data Fig. 1a**) and flow cytometry (**Extended Data Fig. 1b**), the SARS-CoV-2 susceptible cell lines, such as HK-2, Calu-3, HepG2, Huh-7.5, and Caco-2, showed detectable levels of ACE2, both in lysates and on the cell surface, although these levels were substantially lower than those found in the highly susceptible Vero E6 cells. Human vascular endothelial HUVEC cells had measurable ACE2 expression, yet they did not support virus infection under the conditions tested, suggesting that they might lack an important pro-viral factor(s). In contrast, HuH-6 and HuH-7 cells were susceptible to infection, yet lacked detectable levels of ACE2. Nonetheless, overall, the cell susceptibility to SARS-CoV-2 mostly correlated with ACE2 expression (**Fig. 1c**). Our attempts to measure TMPRSS2, another viral entry factor, by Western blot and flow cytometry did not yield reliable results with any of the antibodies tested (data not shown).

### ACE2 and TMPRSS2 expression allows efficient SARS-CoV-2 infection of human cells

We next investigated whether exogenous expression of ACE2 and/or TMPRSS2 in human cells alter their susceptibility to SARS-CoV-2. The cell lines overexpressing ACE2 allowed varying levels of infection whereas TMPRSS2 had mostly no or only subtle effect (**Fig. 2a and Extended Data Fig. 2a**), when entry was probed using vesicular stomatitis virus (VSV) pseudoparticles carrying the SARS-CoV-2 spike protein and expressing the green fluorescent protein (GFP)^14^. However, when ACE2 and TMPRSS2 were expressed together, most of the cell lines became highly susceptible to infection, with 293T, Calu-6, HuH-6, Huh-7.5, and Caco-2 cells demonstrating considerably higher infection efficiency than Vero E6 cells. Human intestinal HT29 cells remained refractory to infection even after ACE2/TMPRSS2 co-expression, and HUVEC, HeLa, RD, and MRC5 cells allowed only moderate infection. All cell lines were highly permissive to wild-type VSV replication (**Extended Data Fig. 2b**), suggesting that S protein-mediated entry was the major factor determining cell susceptibility.

**Fig. 2:**
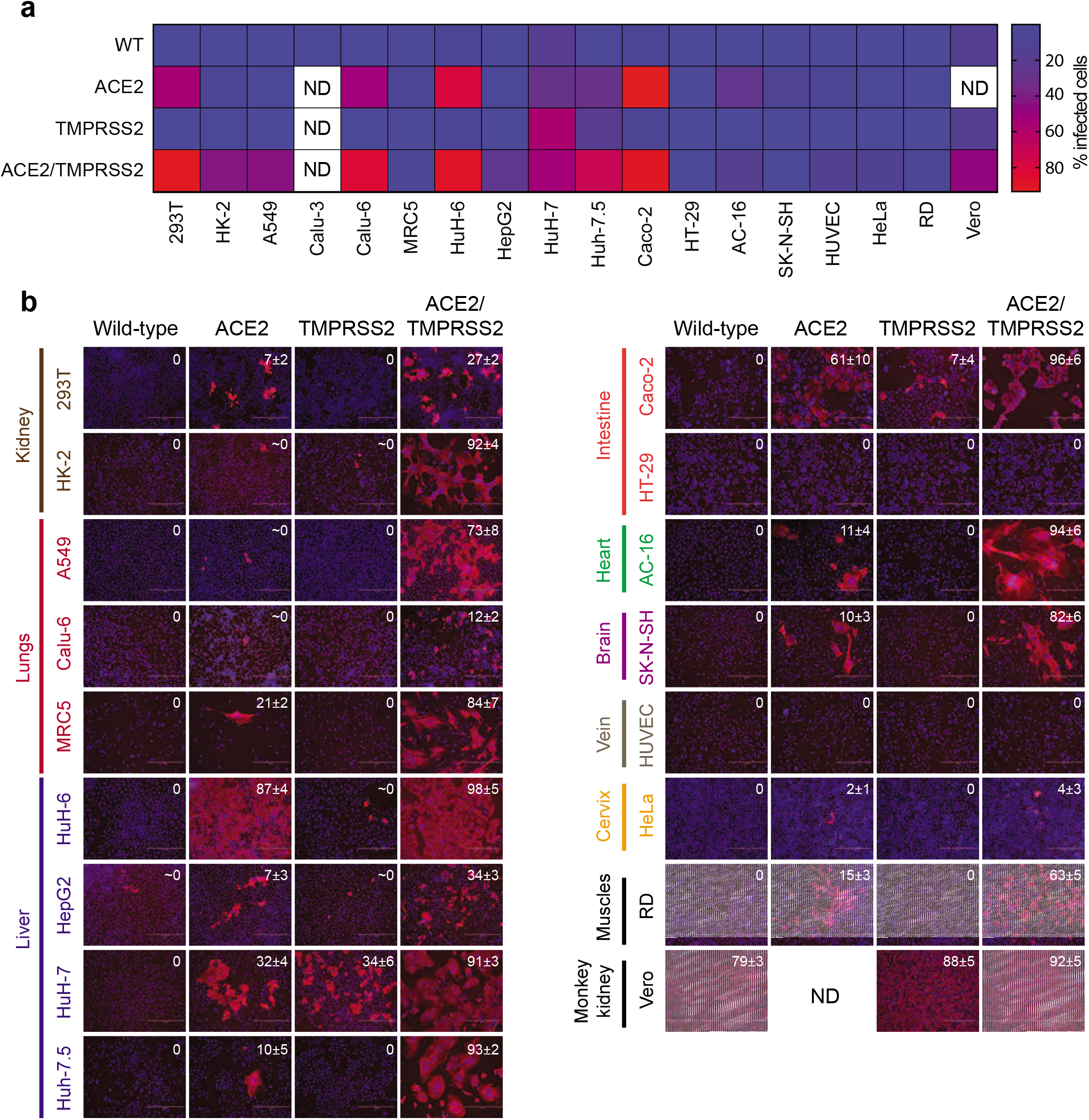
Expression of ACE2 and TMPRSS2 synergistically enhanced SARS-CoV-2 infection of multiple cell lines. **a,** The cells were infected for 24h with VSV pseudotyped with SARS-CoV-2 spike and the infection efficiency was monitored by flow cytometry for GFP, encoded by the VSV genome. The mean ± standard deviation of triplicate samples is plotted. ND, not determined. **b**, The cells were infected for 24h with SARS-CoV-2 at an MOI of 0.01 followed by IF analysis of the viral N-protein (red). The nuclei were stained with DAPI. The mean percentage of positive cells ± standard deviation of three biological replicates is shown (n = 3). ND, not determined.

This pattern was confirmed with authentic SARS-CoV-2. We infected the cells at a low MOI of 0.01 and identified the infected cells by IF at 24 hpi. Consistent with the pseudoparticle assay, ACE2 overexpression allowed a varying degree of infection in several cell lines, with the number of positive cells ranging between 2-87% (**Fig. 2b**). The most remarkable infection efficiency was observed when ACE2 and TMPRSS2 were expressed together, with nine of the 17 human cell lines achieving over 70% positivity, numbers comparable to Vero E6 cells. Interestingly, in congruence with the pseudoparticle assay, HT29 cells remained resistant to infection even after ACE2/TMPRSS2 co-expression (**Fig. 2b**). Overall, these results indicate that exogenous expression of ACE2 and TMPRSS2 renders most of the human cell lines highly susceptible to SARS-CoV-2 infection.

### Proteomic analysis of human cell lines infected with SARS-CoV-2

The development of human cell lines that showed uniform infection allowed us to employ global proteomic analysis to assess the impact of SARS-CoV-2 on host cells without the complication of disambiguating a mixture of uninfected and infected cells. The ACE2/TMPRSS2-expressing A549 (lung), Caco-2 (intestine), HuH-6 (liver), AC-16 (heart), SK-N-SH (brain), and HK-2 (kidney) cells were infected with SARS-CoV-2 at an MOI of 1 and subjected to global proteomic analysis at two different times post-infection (12 and 24 hpi for A549, Caco-2, and HuH-6; 8 and 12 hpi for AC-16, SK-N-SH, and HK-2) (**Fig. 3a**). To achieve a synchronized infection, we adsorbed the virus onto cells on ice for 1h followed by incubation at 37°C. The infections were performed in triplicate, and in-parallel analyses of time-matched, mock-infected cells were performed for comparison. Depending on the cell type, 50-80% cells were found infected at the first time of harvest and 80-100% at the second time (**Fig. 3b, top two panels**).

**Fig. 3:**
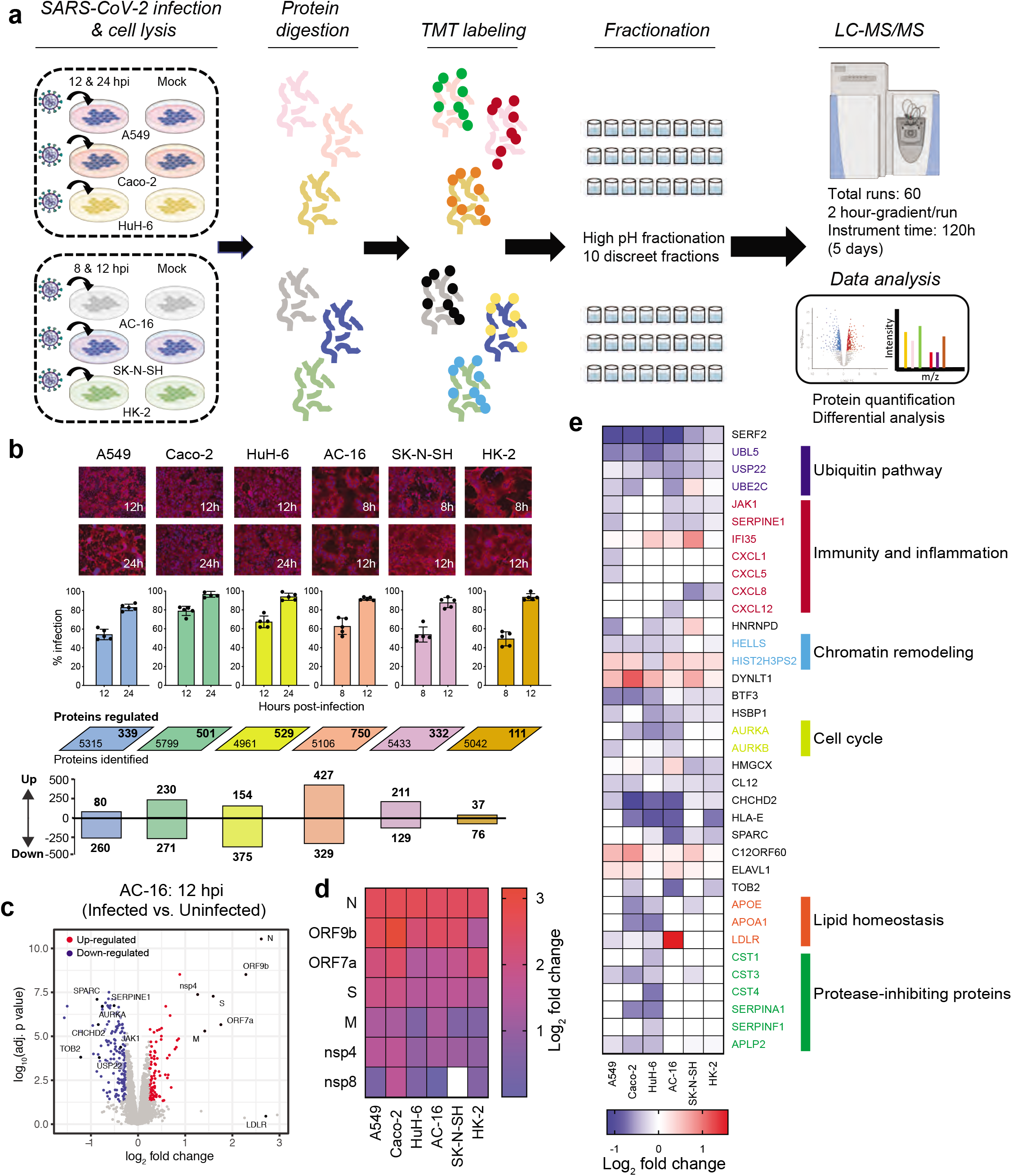
Global proteomic analysis of SARS-CoV-2-infected cells revealed several differentially regulated proteins. **a,** Schematics of the proteomics pipeline. Total protein was extracted from the SARS-CoV-2-infected and uninfected cells, trypsinized, and isotope (TMT) labeled. The peptides for each cell line were separately pooled, fractionated, sequenced, and quantified by LC-MS/MS. **b**, The cells were infected with SARS-CoV-2 at an MOI of 1 and processed at 12h and 24 h (A549, Caco-2, and HuH-6) or at 8h and 12h (AC-16 and SK-N-SH) for IF and proteomic analysis. The top panel shows the IF images (red color: viral N protein; blue color: DAPI). The percentage of positive cells was measured and plotted as a mean ± standard deviation of five microscopic fields (n = 5). The rhomboids show the total number of proteins identified in uninfected and infected cells (lower left corner) as well as the total number of regulated proteins in each cell line across both time points following SARS-CoV-2 infection (upper right corner). Numbers of distinct proteins up- or down-regulated after infection are shown in the bottom panel. In certain cases, the total numbers of up- and down-regulated proteins do not match with the numbers shown in rhomboids. This is due to some proteins downregulated at one time point and upregulated at the other time point. **c**, Volcano plot of proteins regulated in AC-16 cells upon SARS-CoV-2 infection. Proteins enriched in infected cells are shown in red, while those depleted in blue. Black color is used for the proteins labeled with their names. **d**, Heat map showing the abundance of viral proteins in different cell lines. **e**, Heatmap visualization of cellular proteins found to be differentially regulated in more than one cell line. The pathways to which some of these proteins belong are shown on the right.

Whole proteomic analysis identified around 5,000 proteins in both uninfected and infected cells across cell lines. Despite the fact that MS^2^-based TMT ratios are heavily affected by ratio compression^15^, the high reproducibility among replicates allowed us to identify hundreds of differentially regulated proteins in each cell line (**Fig. 3b, middle and bottom panels**). As expected, the most highly enriched candidates in the infected cells were the viral proteins (**Fig. 3c and Extended Data Fig. 3**), including structural proteins, such as, spike (S), membrane (M), and nucleocapsid (N); accessory proteins, such as, ORF7a and ORF9b; and non-structural components mapping to the polyprotein PP1ab (**Fig. 3d**), which is consistent with earlier studies^16^.

Several cellular proteins were found to be differentially affected by SARS-CoV-2 in diverse or distinct cell types (**Supplementary Information and Extended Data Fig. 4**). We mainly focused our analysis on proteins broadly targeted by the virus across cell lines (**Fig. 3e**). Among such proteins were the members of the ubiquitin pathway, such as USP22, UBL5, and UBE2C, which were downregulated in several cell lines. Similarly, HNRNPD (aka AUF1), a protein known to inhibit enteroviruses through degradation of the viral RNA^17^, was depleted in SARS-CoV-2-infected cells. Several chemokines, which are essential mediators of inflammation and play important roles in controlling viral infections, such as CXCL1, CXCL5, CXCL8, and CXCL12 were also among the downregulated proteins. Consistent with previous reports, several proteins involved in the cell cycle regulation, including AURKA and AURKB, were diminished upon SARS-CoV-2 infection^18^. We also identified differential regulation of several innate immune components in the proteomic dataset. For instance, JAK1 and SERPINE1 were downregulated, whereas IFI35, which has been shown to negatively regulate antiviral responses, was enriched following SARS-CoV-2 infection (**Fig. 3e**).

We performed Western blot analysis on a subset of proteins that showed differential regulation in our proteomics analysis. As controls, we included two additional viruses, yellow fever virus (YFV) and Coxsackievirus B3 (CVB3), to rule out any general stress responses induced upon a positive-sense RNA virus infection (**Fig. 4a**). Western blotting largely revealed the protein expression patterns consistent with the proteomics results (**Fig. 4b**). As an example, USP22 was depleted in all cell lines tested, while APOE was impaired in a small subset of cells. For the most part, YFV and CVB3 infection did not affect the selected proteins, with the exception of TOB2, that was depleted in all virus-infected cells. These results confirmed the authenticity of our proteomic dataset and indicate that most of the alterations we detected in the infected cells were specifically due to SARS-CoV-2 infection rather than simply a generic response to viral invasion.

**Fig. 4:**
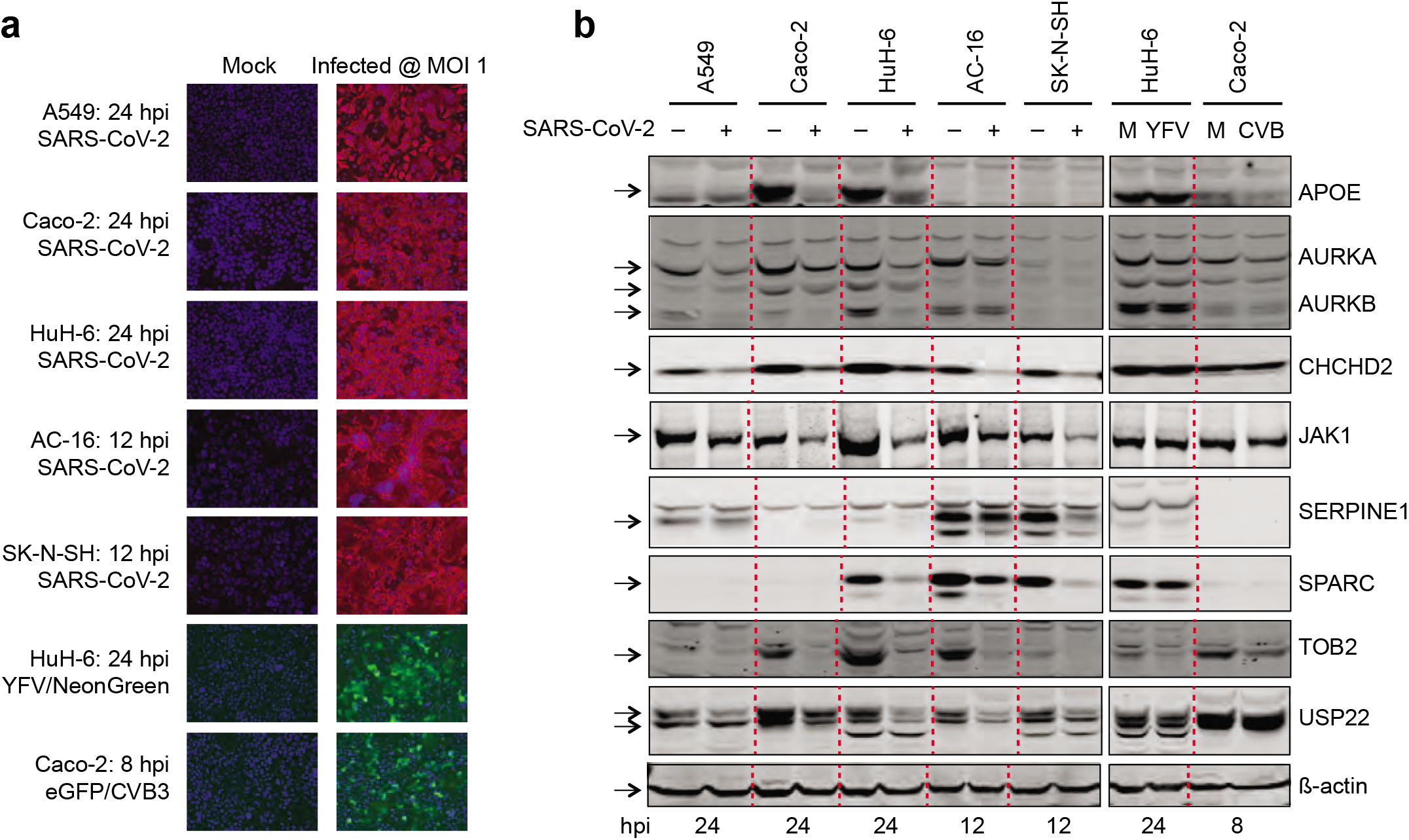
Western blot confirmed the proteomic results. **a**, ACE2/TMPRSS2-expressing A549, Caco-2, HuH-6, AC-16, and SK-N-SH cells were infected with freshly prepared SARS-CoV-2 at an MOI of 1. As a control, HuH-6 cells were infected with YFV 17D virus containing the NeonGreen reporter (MOI of 1) and Caco-2 cells with CVB3 containing the GFP reporter. The cells were fixed at the indicated times and processed for IF and/or imaging. **b**, The cells were lysed in a RIPA buffer followed by the Western blot analysis of the indicated proteins. An equal amount of total protein (25 *μ*g), as quantified by the BCA assay, was loaded in each lane. The black arrows indicate the protein bands of expected sizes. M, Mock.

### SARS-CoV-2 blocks the interferon signaling pathway

Among the cellular proteins downregulated across all cell lines was Janus kinase 1 (JAK1) (**Fig. 4b**), a key signaling molecule downstream of interferons (IFN) and other cytokines, such as interleukin (IL)-2, IL-4, IL-6, and IL-7^19,20^. This prompted us to examine other components of the IFN signaling pathway. Tyrosine kinase 2 (TYK2) showed decreased expression similar to JAK1 and was downregulated in most of the cell lines tested (**Fig. 5a**). Janus kinase 2 (JAK2) exhibited cell type-dependent inhibition; it was diminished in A549, Caco-2, and HuH-6 cells, but not in AC-16 and SK-N-SH cells. When we tested the abundance of type I IFN receptors, we found strong inhibition of IFNAR1, but not IFNAR2, following SARS-CoV-2 infection (**Fig. 5a**). Detailed time course investigation showed that most of the JAK-STAT components were depleted early in infection, indicating a quick active repression of their abundance as opposed to collateral effect of generalized gene inhibition often observed late during infection (**Fig. 5b and Extended Data Fig. 5a**).

**Fig. 5:**
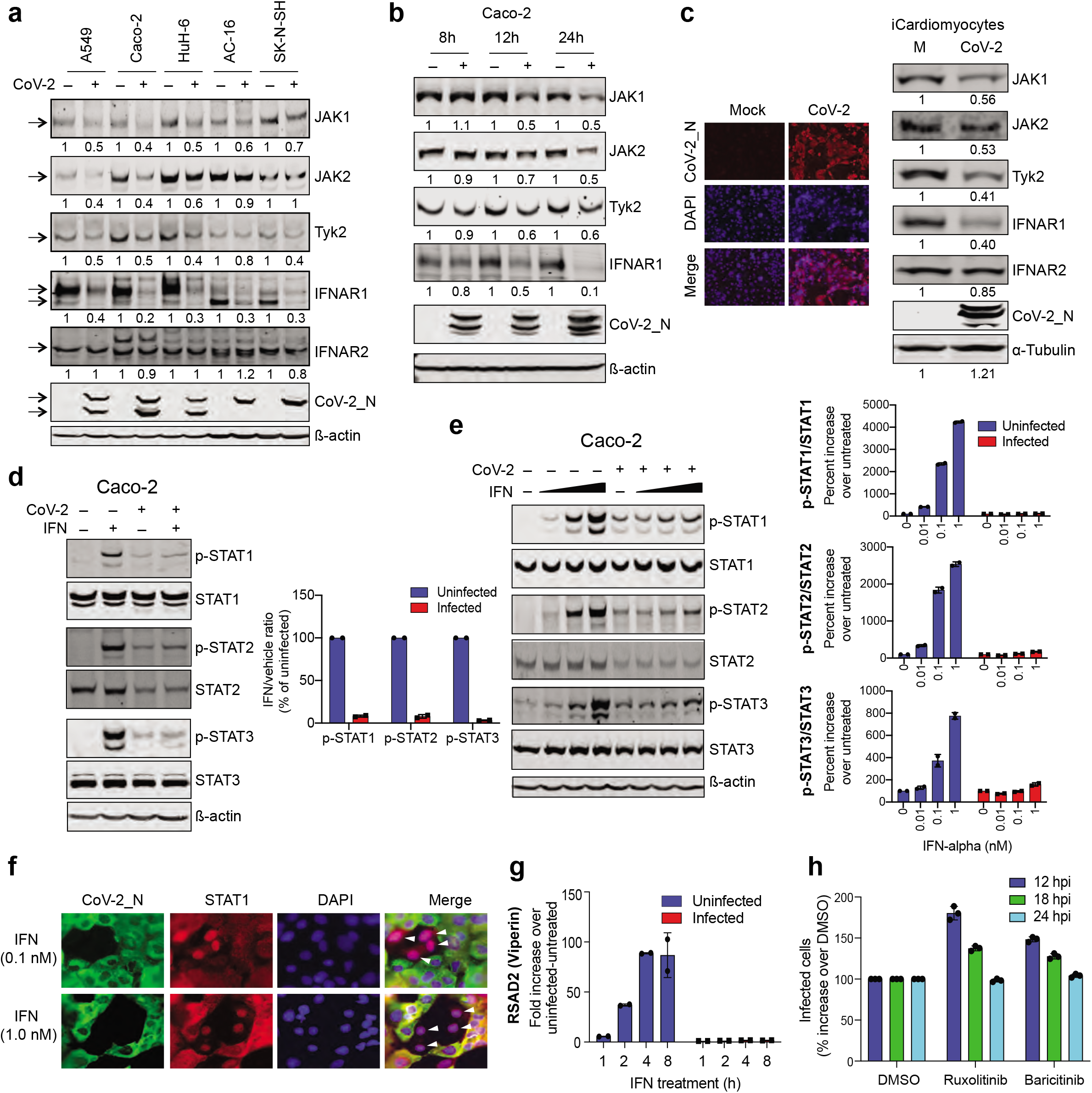
SARS-CoV-2 inhibited IFN signaling. **a**, ACE2/TMPRSS2-expressing A549, Caco-2, and HuH-6 cells were infected with SARS-CoV-2 for 24h, and AC-16 and SK-N-SH for 12h followed by Western blot. The numbers indicate the band intensities, with the uninfected cells arbitrarily set at 1. The black arrows indicate the protein bands of expected sizes. **b**, Caco-2 cells were infected with SARS-CoV-2 at an MOI of 1 and harvested at 8, 12, and 24h post-infection followed by detection of the indicated proteins by Western blot. The band intensities relative to uninfected cells are shown. **c**, hiPSC-CMs were infected with SARS-CoV-2 at an MOI of 5 for 72h, followed by IF (left panel) and Western blot (right panel). The relative band intensities are shown. **d**, Uninfected Caco-2 cells or the ones infected with SARS-CoV-2 (MOI of 1) for 24h were treated with human IFN alpha-2a (1 nM) or, as a negative control, with vehicle (PBS) for 30 min, followed by Western blot. The band intensities of the phospho-STATs were normalized against the total STATs and plotted in the right panel as a percentage of uninfected cells. The data are presented as mean ± standard deviation of two independent Western blots (n = 2). **e**, Uninfected or SARS-CoV-2-infected Caco-2 cells (24 hpi) were treated with 0, 0.01, 0.1, and 1 nM of IFN alpha-2a for 30 min and subjected to Western blot. The band intensities from two independent Western blots, calculated as in **d**, are presented as mean ± standard deviation in the right panel. The intensities for untreated cells were set at 100, and the percent increase in IFN-treated cells was measured by calculating the ratio between IFN-treated and untreated cells (n = 2). **f**, Caco-2 cells infected with SARS-CoV-2 for 24h were exposed to 0.1 or 1 nM IFN alpha-2a for 30 min and stained for the viral N protein (green) and STAT1 (red). The nuclei were stained with DAPI (blue). The nuclear translocation of STAT1 is indicated with white arrowheads. **g**, Caco-2 cells, uninfected or infected with SARS-CoV-2 for 24h, were treated with IFN (1 nM) for 1, 2, 4, or 8h and the RNA levels of Viperin was measured by RT-qPCR. The values were normalized against RPS11 that served as a housekeeping gene. The data are plotted as mean ± standard deviation of three biological replicates (n = 3). **h,** hiPSC-CMs were infected with SARS-CoV-2 at an MOI of 5 in the presence of DMSO or 5 *μ*M compounds. IF was performed at 12, 18, and 24 hpi and the number of positive cells were counted by Muvicyte (see methods). The data are plotted as mean ± standard deviation of three biological replicates (n = 3).

To confirm the authenticity of these findings in a more physiologically relevant setting, we established a human induced pluripotent cell (hiPSC)-derived model of SARS-CoV-2 infection and monitored the ability of the virus to impair IFN-related proteins. We differentiated hiPSC into cardiomyocytes (hiPSC-CM) and first assessed their ability to support virus infection. IF analysis showed that these cells were highly permissive to SARS-CoV-2 infection, and when infected at an MOI of 5, most of the cells became positive by 72 hpi (**Fig. 5c, left panel**). In these infected cardiomyocytes, we observed strong depletion of JAK1, TYK2, and IFNAR1, but not of IFNAR2, consistent with the cell line data (**Fig. 5c, right panel**). These results clearly indicate that SARS-CoV-2 causes bona-fide inhibition of upstream signaling molecules in the interferon response pathway.

Engagement of IFN to its receptor complex and the subsequent activation of receptor-associated JAK kinases leads to tyrosine phosphorylation, dimerization, and activation of STAT proteins^21^. Because SARS-CoV-2 inhibited the expression of IFN receptor and its associated JAKs, we next determined if this inhibition desensitized the infected cells to IFN treatment. For this, we treated uninfected and infected cells with IFN-α and examined the phosphorylation of three main STATs involved in IFN signaling, namely STAT1,2, and 3. As expected, IFN treatment of uninfected cells for 30 minutes caused extensive phosphorylation of all three STATs without influencing the abundance of total proteins (**Fig. 5d and Extended Data Fig. 5b**). In contrast, however, the virus-infected cells were highly resistant to IFN, with some cell types showing no change in STAT phosphorylation while others registering only subtle effects. The strongest effect was seen in Caco-2 cells, where STAT phosphorylation was inhibited by over 90% following virus infection (**Fig. 5d, right panel**). Interestingly, the abundance of total STAT2 protein was also diminished in infected cells, indicating an additional layer of virus-imposed inhibition of IFN signaling (**Fig. 5d**). To further confirm these findings, we performed a dose-response analysis of STAT phosphorylation upon IFN treatment. While escalating doses of IFN caused increasing phosphorylation of STATs in uninfected cells, only negligible effect was seen in SARS-CoV-2-infected cells (**Fig. 5e**).

Once activated by phosphorylation, STAT proteins make homo-or heterodimers and translocate to the nucleus where they activate the expression of interferon stimulated genes (ISGs)^21^. Consistent with the decreased phosphorylation of STAT proteins following SARS-CoV-2 infection, we observed strong inhibition of STAT1 nuclear translocation. While most of the STAT1 protein in uninfected cells moved to the nucleus within 30 min of IFN treatment, it largely remained in the cytoplasm in SARS-CoV-2-infected cells (**Fig. 5f**). This was associated with a significant loss of ISG induction. We treated uninfected and virus-infected cells with IFN for 1, 2, 4, and 8h and examined the expression of two ISGs, ISG15 and RSAD2 (a.k.a. Viperin) by RT-qPCR. Remarkably, the expression of both ISGs was completely blocked in the infected cells as compared to their uninfected counterparts that showed between 3- and 85-fold upregulation of ISGs following IFN treatment (**Fig. 5g and Extended Data Fig. 6**). In line with these findings, chemical inhibition of the JAK-STAT pathway with JAK inhibitors, Ruxolitinib and Barcitinib, increased SARS-CoV-2 infection of hiPSC-CMs, as evidenced by 1.5-to 2-fold increase in the percentage of virus-infected cells at 12 hpi, although this replicative advantage was lost by 24 hpi (**Fig. 5h**). In all, these results indicate that SARS-CoV-2 targets multiple components of the JAK-STAT axis and transforms infected cells into an interferon non-responsive state (**Fig. 6**).

**Fig. 6:**
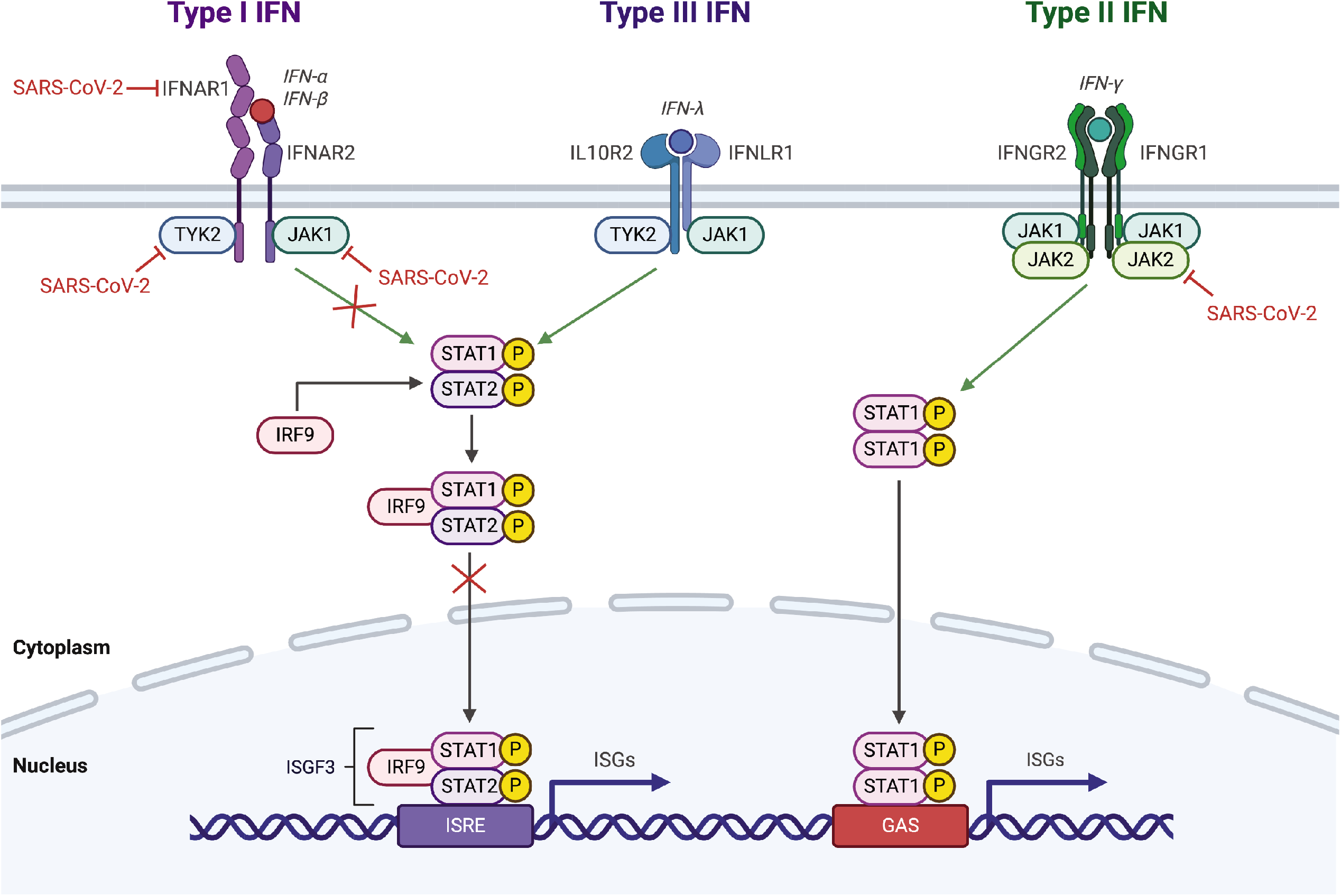
Model of the JAK-STAT inhibition by SARS-CoV-2. Engagement of IFNs to their receptors causes receptor dimerization, bringing JAKs attached to the cytoplasmic side of the receptors into close proximity. The JAKs then phosphorylate each other through a process called transphosphorylation, leading to their activation. The activated JAKs phosphorylate STAT proteins, which then form hetero- or homo-dimers and recruit interferon regulatory factor 9 (IRF9) to make a trimolecular complex, called ISGF3, that translocates to the nucleus to induce expression of hundreds of interferon-stimulated genes (ISGs). These ISGs create an antiviral environment in the cell leading to elimination of the replicating virus and inhibition of new viral invasions. SARS-CoV-2 inhibits the JAK-STAT pathway at multiple steps indicated with red inhibition arcs. The ultimate effect of this inhibition is a blunted antiviral response unable to tackle the virus.

## DISCUSSION

We report an immune evasion strategy that SARS-CoV-2 uses to antagonize IFN signaling and establish infection in diverse tissue types. Being amongst the earliest and most potent of the immune responses, IFN signaling poses an immediate threat to viruses and can quickly eliminate them from the infected cells. Two recent reports have shown that inborn errors in IFN signaling and the presence of anti-IFN auto-antibodies can predispose individuals to a life-threatening COVID-19 disease, highlighting the importance of IFN immunity in SARS-CoV-2 infection^22,23^. Our findings complement these reports and show that SARS-CoV-2 has evolved a suite of mechanisms to counteract the effector functions of interferon. This inhibition of IFN signaling is also expected to attenuate the production of IFN, as these two processes are tightly regulated by a positive feedback loop, with early IFNs inducing the expression of viral sensors^24^, interferon regulatory factor 7 (IRF7), and other signaling molecules^25^, which regulate the expression of IFN^26^. The overall outcome of this inhibition is an attenuated antiviral state supporting uninterrupted viral propagation in disparate tissues.

Several studies are currently evaluating the clinical efficacy of IFN alone or in combination with antiviral compounds for the treatment of COVID-19 (https://clinicaltrials.gov). This is based on the supposition that IFNs can help clear infection in two ways. First, by acting on infected cells and eliminating the replicating virus. Second, by creating an antiviral state in uninfected cells to block viral spread. There is support for the latter mechanism of action from recent studies showing that pre-treatment with IFN can effectively inhibit SARS-CoV-2 replication in cell culture^27,28^. However, our results show that once the viral replication is established inside a cell, its ability to respond to IFN is severely compromised, suggesting that the former potential protective mechanism of IFN treatment may not function in COVID-19 patients. In a clinical setting, by the time a patient receives IFN treatment, a large number of cells are infected with the virus, and these would be expected to be non-responsive to interferon treatment.

Considering this scenario, the timing of IFN therapy relative to infection is expected to be the key determinant of outcome. In a recent retrospective study of 446 patients with COVID-19, early use of IFN was found to be protective, whereas its late administration was associated with delayed recovery and increased mortality^29^. Similarly, the World Health Organization’s large-scale, multi-country clinical trial showed no protective effect of IFN in hospitalized COVID-19 patients with late-stage disease (https://www.medrxiv.org/content/10.1101/2020.10.15.20209817v1). The same has been previously reported in mice infected with a related human coronavirus, MERS-CoV, where early administration of IFN protected the animals from lethal infection, whereas delayed administration of IFN failed to inhibit virus replication^30^.

Of note, the activation of the JAK-STAT pathway has been shown to stimulate the production of IL-6 and other inflammatory cytokines^31^, which in turn attract an army of immune cells to the site of infection to orchestrate the destruction of infected cells. Inhibition of the JAK-ST AT axis by SARS-CoV-2 can therefore suppress the production of cytokines, thereby disrupting a potent and timely inflammatory response. The interferon pathway is also implicated in a crosstalk between innate and adaptive immunity^32^. Loss of bridging between these two immunity arms and the resulting timing mismatch has been postulated to cause severe disease in COVID-19 patients^33,34^. A recent study that examined peripheral blood from patients with COVID-19 of varying severity also reported a strong correlation between impairment of IFN responses and the clinical outcome of infection^35^. These two observations, when combined, firmly point to the possibility of aberrant IFN responses to be at the heart of COVID-19 pathogenesis.

Our study reports a suite of human cell lines suitable for SARS-CoV-2 studies. Exogenous expression of ACE2 has been demonstrated to improve the susceptibility of a few human cell lines, such as A549 and HeLa, to SARS-CoV-2^36,37^, but our results show that efficient infection can only be achieved when ACE2 and TMPRSS2 are co-expressed. This finding is important because it provides a conceptual framework for generating efficient infection models for SARS-CoV-2. Human ACE2-expressing mice that are currently used to investigate SARS-CoV-2^38^ do not recapitulate the full spectrum of the human disease (reviewed in^39^). From our results, it is tempting to speculate that co-expression of human ACE2 and TMPRSS2 might improve the quality and physiological relevance of these mouse models.

The human cell lines we generated represent a rich resource that can be used to investigate cellular pathways broadly targeted by SARS-CoV-2 in diverse tissue types. While immortalized and cancerous cell lines often incompletely recapitulate the genotypic and phenotypic profile of their tissues of origin, many cell lines maintain an important degree of tissue identity^40^. This is reflected in our proteomic dataset. As an example, APOE, a protein regulated upon virus infection and which we validated by Western blot, was only found to be altered in liver and intestinal cell lines, two tissue types known to express this protein to high levels^41^. The results obtained from hiPSC-CMs closely matched with those from the cardiac cell line AC-16, providing further validity to our cell line approach (**Fig. 5a, c**).

Recent studies have reported proteomic profiling of SARS-CoV-2 infection in Caco-2^16^ and A549 cells (https://www.biorxiv.org/content/10.1101/2020.06.17.156455v1). These studies, however, employed relatively suboptimal infection models, and thus relied on a mosaic of infected and uninfected cells. We found little overlap between our results and those reported by others. This could be due to the differences in viral isolates, proteomic platforms, and analyses pipelines, but is likely to also be due to disparities in the number of infected cells in the cell population. Our infection models supported quick, uniform virus infection allowing us to identify SARS-CoV-2-specific changes as opposed to generalized stress responses, as evident from the Western blot analysis of select hits in cells infected with two non-coronaviruses (**Fig. 4b**).

In all, this study uncovers the phenomenon of SARS-CoV-2-mediated desensitization to IFN. These are critical findings that will enhance our understanding of how SARS-CoV-2 mutes the innate immune system to sustain its replication, leading to impaired adaptive immune responses and consequently severe disease. The knowledge gained from these studies will provide the foundation for the development of therapeutic interventions aimed at boosting cell-intrinsic antiviral responses and inform the design of attenuated viruses as potential vaccine candidates.

## METHODS

### Cells, antibodies, and chemicals

All cell lines were incubated at 37°C and 5% CO_2_ in a humidified incubator. Human embryonic kidney HEK293T cells (ATCC; CRL-3216), human cervical carcinoma HeLa cells (ATCC; CCL-2), human colorectal adenocarcinoma HT-29 cells (ATCC; HTB-38), human hepatoblastoma HuH-6 cells (JCRB-0401), human hepatocellular carcinoma HuH-7 cells (JCRB-0403) and its derivative Huh-7.5 cells, human hepatocellular carcinoma HepG2 cells (ATCC; HB-8065), human lung anaplastic carcinoma Calu-6 cells (ATCC; HTB-56), human lung adenocarcinoma A549 cells (ATCC; CCL-185), human normal lung fibroblast MRC-5 cells (ATCC; CCL-171), human kidney papilloma HK-2 cells (ATCC; CRL-2190), human neuroblastoma SK-N-SH cells (ATCC; HTB-11), African green monkey kidney Vero E6 cells, and human muscle rhabdomyosarcoma RD cells (ATCC; CCL-136) were maintained in DMEM (Gibco; #11995-065) containing 10% FBS and 1X non-essential amino acids, human colorectal adenocarcinoma Caco-2 cells (ATCC; HTB-37) were maintained in the same medium but containing 20% FBS, human cardiomyocyte AC16 cells (Millipore; SCC109) in DMEM/F12 (Gibco; #11330-032) containing 12.5% FBS, human lung adenocarcinoma Calu-3 cells (ATCC; HTB-55) in MEM (Gibco; #11095-080) containing 10% FBS, and human umbilical vein endothelial HUV-EC-C cells (ATCC; CRL-1730) in RPMI 1640 medium (Gibco; #11875-093) containing 10% FBS. Mycoplasma negative status of all cell lines was confirmed. Cells stably expressing human ACE2 and TMPRSS2 were generated by lentiviral transduction followed by selection with appropriate selection drugs.

Anti-SARS-CoV nucleocapsid (N) protein antibody (Rockland; #200-401-A50) was used for detection of SARS-CoV-2 N protein by IF. Anti-hACE2 antibodies included rabbit monoclonal antibody EPR4435(2) (Abcam; #ab108252) for Western blot and goat polyclonal antibody (R&D Systems; #AF933) for flow cytometry. Goat IgG isotype control antibody was from Invitrogen (#02-6202) and Anti-VSV-G antibody from Kerafast (#EB0010). Antibodies for validation of our proteomics results and the follow-up studies included APOE (Proteintech; #18254-1-AP), AURKA (Bethyl Laboratories; #A300-071A), CHCHD2 (Proteintech; #19424-1-AP), JAK1 (Santa Cruz Biotechnology; #sc-277), LDLR (Bethyl Laboratories; #A304-417A), SERPINE1 (Novus Biologicals; NBP1-19773), SPARC (Proteintech; #15274-1-AP), TOB2 (Proteintech; #13607-1-AP), USP22 (Novus Biologicals; #NBP1-49644), JAK2 (Proteintech; #17670-1-AP), TYK2 (Proteintech; #67411-1-Ig), JAK 3 (Cell Signaling Technologies; #3775), p-STAT1 (Cell Signaling Technologies; #9171), STAT1 (Cell Signaling Technologies; #14994), pSTAT-2 (Cell Signaling Technologies; #4441), STAT2 (Santa Cruz Biotechnology; #sc-476), pSTAT3 (Cell Signaling Technologies; #9145), STAT3 (Santa Cruz Biotechnology; #sc-482), β-actin (Invitrogen; #AM4302), and α-Tubulin (ECM Biosciences; #TM4111).

Ruxolotinib (INCB018424) was from Selleck Chemicals (#S1378) and Baricitinib phosphate from Fisher Scientific (#50-201-3519).

### Plasmids

Expression plasmid encoding the spike protein of SARS-CoV-2, pCG1_SARS-2_S, has recently been described and was a kind gift from Stefan Pohlmann^42^. The plasmids pCAGGS_TMPRSS2-Flag and pcDNA3.1_ACE2, encoding human ACE2 and TMPRSS2, respectively, were obtained from Thomas Gallagher (Loyola University). We transferred ACE2 and untagged TMPRSS2 to the lentiviral pLOC vector to obtain pLOC_hACE2_PuroR and pLOC_hTMPRSS2_BlastR, respectively.

### SARS-CoV-2 stock preparation and titration

We used 2019-nCoV/USA-WA1/2020 isolate (NCBI accession number: MN985325) of SARS-CoV-2, obtained from the Centers for Disease Control and Prevention and BEI Resources. To generate the passage 1 (P1) virus stock, we infected Vero E6 cells, seeded one day prior into a 175 cm^2^ flask at a density of 10 million cells, with the master stock diluted in 10 ml of Opti-MEM. Following virus adsorption to the cells at 37°C for 1h, we added 15 ml of DMEM containing 10% FBS and 1X penicillin/streptomycin. The next day, we removed the inoculum, rinsed the cell monolayer with 1X PBS, and added 25 ml of fresh DMEM containing 2% FBS. Two days later, when the cytopathic effect of the virus was clearly visible, as evidenced by a large number of round floating cells, we collected the culture medium, passed it through a 0.2*μ* filter, and stored at −80°C. We then prepared the P2 working stock of the virus by infecting Vero E6 cells with the P1 stock at a multiplicity of infection (MOI) of 0.1 plaque forming units (PFU)/cell and harvesting the culture medium three days later.

To determine the titer of our viral stock by plaque assay, we seeded Vero E6 cells into a 12-well plate at a density of 2.5 x 10^5^ cells per well. The next day, the cells were infected with serial 10-fold dilutions of the virus stock for 1 h at 37°C. We then added 1 ml per well of the overlay medium containing 2X DMEM (Gibco; #12800017) supplemented with 4% FBS and mixed at a 1:1 ratio with 1.2% Avicel (DuPont; RC-581) to obtain the final concentrations of 2% and 0.6% for FBS and Avicel, respectively. Three days later, the overlay medium was removed, the cell monolayer was washed with 1X PBS and fixed for 30 minutes at room temperature with 4% paraformaldehyde. Fixed cells were then washed with 1X PBS and stained for 1h at room temperature with 0.1% crystal violet prepared in 10% ethanol/water. After rinsing with tap water, the number of plaques were counted and the virus titer was calculated. The titer of our P2 virus stock was 1 x 10^7^ PFU/ml.

### VSV pseudoparticle production and infection

VSV pseudoparticles carrying the SARS-CoV-2 S protein were generated as previously described^42^. Briefly, 293T cells were seeded into a 6-well plate at the density of 5×10^5^ cells per well. The next day, the cells were transfected with 6 *μ*g of pCG1_SARS-2_S using X-tremeGENE 9 DNA transfection reagent (Sigma-Aldrich). Twenty hours later, the cells were inoculated with a replication-competent vesicular stomatitis virus (VSV) modified to contain an expression cassette for GFP in place of the VSV-G open reading frame, VSV*ΔG-GFP (kindly provided by Stefan Pohlmann of the Leibniz Institute for Primate Research (DPZ), Gottingen, Germany). Following 2h incubation of cells with VSV*ΔG-GFP at 37°C, the inoculum was removed and the cells were washed three times with FBS-free DMEM. Two ml of DMEM/10%FBS supplemented with anti VSV-G antibody was then added to each well to neutralize any residual input VSV. Pseudoparticles were harvested 20h later, passed through 0.45 *μ* filter, and stored at −80°C.

For infection experiments, target cells grown in 24-well plates to the confluency of 70-80% were infected with pseudoparticles. Twenty-four hours later, the cells, including those floating around in the culture medium due to cytopathic effects of VSV replication, were collected and fixed in 4% PFA. Flow cytometry was performed to count the percentage of GFP positive cells.

### Quantitative real-time PCR (RT-qPCR)

To measure the mRNA abundance of ISGs, total RNA was isolated from cells using Qiagen RNeasy Plus Mini Kit (Qiagen #74134), according to the manufacturer’s instructions, with an additional on-column DNase treatment (Qiagen; #79256). RT-qPCR was performed using Luna^®^ Universal One-Step RT-qPCR kit (New England Biolabs; #E3005L). Briefly, a 12 *μ*l of reaction mixture containing 2 *μ*l of RNA, 0.4 *μ*M of each forward and reverse primer, 0.6 *μ*l of the 20X Luna WarmStart^®^ RT Enzyme Mix, and 6 *μ*l of the 2X Luna Universal One-Step Reaction Mix was subjected to one-step RT-qPCR using Applied Biosystems QuantStudio 3 (ThermoFisher Scientific), with the following cycling conditions; reverse transcription at 50°C for 10 min, initial denaturation at 95°C for 2 min followed by 40 cycles of denaturation at 95°C for 15 sec and annealing/extension at 60°C for 1 min, ending with the melt curve analysis of the PCR product from 65°C to 95°C, rising in 0.5°C per second increments, waiting for 30 sec at 65°C and for 5 sec at each step thereafter, and acquiring fluorescence at each temperature increment. The Cq values were determined using the QuantStudio™ Design and Analysis software V1.5.1. RPS11 was used as a housekeeping gene and the ISG Cq values were normalized against this gene to calculate the fold change between untreated and IFN-treated cells.

### Immunofluorescence

Virus-infected cells were fixed in 4% paraformaldehyde for 30 minutes. The fixative was removed and the cell monolayer washed twice with 1X PBS. The cells were then permeabilized and incubated overnight at 4°C with anti-SARS-CoV Nucleocapsid antibody (1:2,000 dilution). The cells were then washed 5 times with 1X PBS and stained with Alexa Fluor 568-conjugated goat anti-rabbit secondary antibody (1:1000 dilution) (Invitrogen; #A11008) in the dark at room temperature for 1h and counterstained with DAPI. Images were captured using EVOS M5000 Imaging System (ThermoFisher Scientific). For quantitative analysis of the fixed cell images, we used the MuviCyte Live-Cell Imaging System (PerkinElmer, Waltham, MA). We acquired images of multiple microscopic fields per well using a 10X objective lens and counted the number of DAPI- and viral antigen-positive cells. For each of those images, we then calculated the percentage of DAPI-positive cells expressing the viral antigen and plotted the mean±SD of multiple images for each condition.

### Flow cytometry

For cell surface analysis of ACE2, we harvested cells and washed them in FACS Buffer (2% FBS in 1X PBS). Cells were resuspended in 1:50 dilution of human F_C_ blocking solution (BioLegend; #422302) and incubated on ice for 10 min. Human ACE2 antibody or goat IgG isotype control was then added to the cells to obtain the final concentration of 5 μg/mL followed by 1h incubation on ice. The cells were washed with FACS buffer and incubated for 30 minutes on ice in the dark with 1:400 dilution of Alexa Fluor 488 donkey anti-goat secondary antibody (Invitrogen; #A11055). The cells were washed and resuspended in FACS buffer. Data were collected using a BD LSR II flow cytometer and analyzed with FlowJo software (version 10).

### Sample preparation for proteomic analysis

We infected cells grown to the confluency of 90-95% in 6-well plates with SARS-CoV-2 at an MOI of 1. To synchronize virus entry into cells, the infection was carried out in a small volume of Opti-MEM (400 *μ*l per well) on ice for 1 h, followed by removal of the virus inoculum, addition of 2 ml of DMEM containing 2% FBS, and incubation at 37°C. To harvest cells, the culture medium was removed and the cell monolayer was washed twice with 1X PBS, followed by cell scraping in a lysis buffer comprising 6M GuHCl, 100 mM Tris pH 8.0, 40 mM chloroacetamide and 10 mM TCEP, supplemented with Complete Mini protease (Roche; #11836170001) and phosphatase inhibitor cocktails (Roche; #04906837001). Lysates were boiled at 100°C for 15 min and sonicated briefly. Total proteins were quantified via Bradford’s assay and normalized prior to digestion with Trypsin (Pierce) at 1:50 ratio (enzyme to protein, w/w) overnight at 37°C. Digestion was quenched with trifluoracetic acid and the peptides were desalted using Sep-Pak C18 columns (Waters Corporation; WAT054955). Desalted peptides were labeled with Tandem Mass Tags (TMT) using TMTPro-16plex isobaric tags (ThermoFisher Scientific; #A44520) as per manufacturer’s instructions. Replicate samples corresponding to each time point for each cell line were labelled separately. The TMT labelled peptides for the respective cell lines were then pooled, desalted, and fractionated via basic reversed-phase fractionation using a mobile phase comprising 0.1% NH4OH and varying acetonitrile concentrations (5%, 10%, 12.5%, 15%, 17.5%, 20%, 22.5%, 25%, 30%, 35%, 40% and 60%). For each cell line, two early (5% and 10%) fractions were orthogonally concatenated with two late (40% and 60%) fractions resulting in total of 10 fractions per cell line for analyses. All fractions were dried in SpeedVac Vacuum Concentrator (ThermoFisher Scientific).

### Mass spectrometry analysis

Dried samples were reconstituted in mobile phase A solvent (2% acetonitrile and 0.1% formic acid) for analysis on the Q-Exactive™ HF-X mass spectrometer (ThermoFisher Scientific), interfaced to the Easy nanoLC1200 HPLC system (ThermoFisher Scientific). The peptides were loaded on a reversed-phase nano-trap column in mobile phase A (75μm i.d. × 2 cm, Acclaim PepMap100 C18 3μm, 100Å; ThermoFisher Scientific; # 164946) and separated over an EASY-Spray column, (ThermoFisher Scientific; #ES803A) using a gradient (6% to 19% over 58 min, then 19% to 36% over 34 min) of mobile phase B (0.1% formic acid, 80% acetonitrile) at a flow rate of 250 nl/min. The mass spectrometer was operated in positive ion mode with a spray voltage of 2100 volts and the data was acquired in a data-dependent acquisition (DDA) mode. Precursor scans were acquired at a resolution of 120,000 FWHM with a maximum injection time of 120 ms. The top 12 abundant ions, with charge states ≥2, were selected for fragmentation by HCD (collision energy 29%) and analyzed at a resolution of 45,000 FWHM with a maximum injection time of 250 ms.

### Analysis of raw mass spectrometry data

All raw data were processed using MaxQuant (Version 1.6.7.0). The acquired tandem spectra were searched against the reference *Homo sapiens* proteome (Taxonomic ID: 9606) FASTA file downloaded from UniProt on April 2017, concatenated with common contaminants and SARS-CoV-2 proteome sequences. TMT reporter ion quantification was performed on MaxQuant using default settings. For searches, cysteine carbamido-methylation was specified as fixed modification and oxidation of methionine and N-terminal protein acetylation were set as variable modifications. Enzyme specificity was set to trypsin and up to two missed cleavages were allowed. The MaxQuant output file designated “ProteinGroups.txt” was used for data normalization and statistical analyses using in-house generated scripts in the R environment.

### Data analysis and pathway enrichment

Bioinformatic analysis was performed using R: A language and environment for Statistical Computing (R Foundation for Statistical Computing, Vienna, Austria. http://www.R-project.org), version 3.6.1. The “ProteinGroups.txt” table corresponding to each cell line was filtered to eliminate entries labelled as reverse hits, potential contaminants, and “only identified by site”. Protein quantitation required at least 70% valid values across all TMT channels. The TMT intensity values were log2 transformed and Loess-normalized. Differentially regulated proteins were defined by implementing a log_2_ fold-change threshold of 0.25 for SARS-CoV-2 vs mock-infected cells. Similar liberal thresholds have been previously described for TMT-based analyses owing to ratio compression^43^. For functional enrichment of proteins based on clustering between the respective time points, Enrichr was used with the Reactome database of pathways. For differential analysis, the LIMMA^44^ R package was used to fit a linear model accounting for the infection versus mock condition at each timepoint. Moderated t-tests were corrected with the Benjamini-Hochberg method for false discovery rate (FDR). Gene set enrichment analysis was performed using the fgsea R package (https://www.biorxiv.org/content/early/2016/06/20/060012) using curated gene libraries^45^ ranked lists where the gene rank is defined as −log(p value)*sign(log2 fold change)^46^. In the case of duplicate proteins mapping to a common gene symbol, the one with the highest absolute value rank was retained.

### Differentiation of hiPSC into cardiomyocytes

Human induced pluripotent stem cells (hiPSCs) from the PGP1 parent line (Coriell Institute; GM23338) engineered to have an endogenous green fluorescent protein tag on one titin allele^47^ were provided by the Seidman Lab. hiPSCs were seeded into tissue culture-treated plates coated with Matrigel (ThermoFisher Scientific; #CB-40230) mixed 1:80 in DMEM/F-12 (ThermoFisher Scientific; #11330-057), maintained in mTeSR1 (StemCell; #85870), and passaged using Accutase (Sigma; #A6964) at 60-90% confluence.

hiPSCs were differentiated into cardiomyocytes (hiPSC-CMs) by small-molecule, monolayer-based manipulation of the Wnt-signaling pathway. Briefly, the cells were grown from day 0 to day 9 in RPMI 1640/GlutaMAX medium (ThermoFisher Scientific; #61870036) containing the insulin-free B-27 supplement (ThermoFisher Scientific; #A1895601). On day 0, the cells were treated with 12 *μ*M CHIR99021 (Tocris; #4423) for 24h to activate WNT signaling. Two days later (on day 3), 5 *μ*M IWP4 (Tocris; #5214) was added to the culture medium for 48h to block the WNT signaling. On day 9, the culture medium was replaced with RPMI 1640/GlutaMAX medium supplemented with insulin-containing B-27 supplement (ThermoFisher Scientific; #17504-044). On day 11, metabolic selection of hiPSC-CMs was started by growing them in a glucose-free RPMI 1640 medium (ThermoFisher Scientific; #11879020) containing 4 mM Sodium DL Lactate solution (Sigma; #L426) for 4 days, replenishing the medium every 2 days.

Following metabolic selection, purified hiPSC-CMs were trypsinized with 0.25% trypsin-EDTA (ThermoFisher Scientific; #25200114) containing 10 *μ*g/ml DNase I (StemCell; #7469) and re-plated into 12-well plates coated with 10 *μ*g/ml human bulk fibronectin (ThermoFisher Scientific; #3560) at a density of 750,000 cells/well. The replating medium comprised RPMI 1640 mixed with the insulin-containing B-27 supplement, 2% fetal bovine serum (Sigma; #F0926), and 5 *μ*M Y-2763 (Tocris: #12543). The cells were maintained in RPMI 1640 medium supplemented with insulin-containing B-27 until day 30-50, with the medium being replenished every two days. For SARS-CoV-2 infection, hiPSC-CMs were seeded into either 96- or 12-well plates coated with 10 *μ*g/ml fibronectin.

### Interferon response assays

To test interferon response in uninfected and SARS-CoV-2-infected cells, we seeded cells either in 6-well plates (for Western blot) at a density of 5 x 10^5^ cells per well, in 12-well plates at a density of 2 x 10^5^ cells per well (for RT-qPCR), or in 96-well plates at a density of 2.5 x 10^4^ cells per well (for IF). The next day, the cells were infected with SARS-CoV-2 at an MOI of 1 or left uninfected. Twenty-four hours later, we treated the cells with different concentrations of IFN alpha-2a for 15-30 min (for Western blot and IF) or for 1, 2, 4, and 8h (for RT-qPCR). The cells were then processed for downstream applications.

### Western blotting

Proteins from various cells were extracted with 1x RIPA buffer containing 1x complete-mini protease inhibitor (Roche; #11836170001) and 1x phosphatase inhibitor cocktail (Roche; #04906837001). Samples were incubated on ice for 30 min and centrifuged at 12,000 xg for 20 min at 4°C. The supernatants were transferred to new ice-cold Eppendorf tubes and protein concentration was measured by the BCA assay using Pierce BCA Protein Assay kit (ThermoFisher Scientific; #23225). Equal amounts of protein were loaded on 4-12% SDS-PAGE gel and transferred onto nitrocellulose membrane. Following staining with primary and secondary (LiCor) antibodies, the bands were visualized by scanning the membrane with the LiCor CLx infrared scanner. The intensity of protein bands was measured in the open source package, ImageJ.

## ACKNOWLEDGEMENT

We thank Drs. Markus Hoffmann and Stefan Pohlmann of Leibniz Institute for Primate Research (DPZ), Gottingen, Germany, for providing the pCG1_SARS-2_S plasmid; Dr. George J. Murphy (Boston University, Boston) for AC-16 cells; Dr. Nader Rahimi (Boston University, Boston) for HUVECs; Dr. Thomas Gallagher (Loyola University, Maywood) for hACE2 and hTMPRSS2 expression plasmids; Dr. Christine E. Seidman (Harvard Medical School, Boston) for iPSCs, and Dr. Elke Muhlberger (Boston University, Boston) for reagents. We also thank Dr. William M. Schneider (The Rockefeller University) and Dr. Mikel Garcia-Marcos (Boston University) for crucial reading of the manuscript. This work was supported by Boston University startup funds (to MS and FD), Evergrande MassCPR awards (to MS and DNK), Peter Paul Career Development Award (to FD), National Cancer Institute (NCI R01CA175382) and National Institutes of Health (NIH R01HL132325) (to VCC), National Science Foundation Graduate Research Fellowship (to JKE), and National Science Foundation Nanosystems Engineering Research Center for Directed Multiscale Assembly of Cellular Metamaterials (to CSC).

## Authors’ contributions

M.S. conceptualized the study. D-Y.C, B.J.C., H.L.C., A.H.T., S.K., D.K., F.D., and M.S. performed the experiments. R.K.G. performed mass spec analyses of SARS-CoV-2-infected cells. J.K.E. and C.S.C. provided hiPSC-CMs. B.B., R.K.G., and M.S. performed data analysis, T.G., N.A.C, V.C.C., D.N.K., and J.H.C. provided reagents and scientific input. M.S., D-Y.C., R.K.G., F.D., and A.E. interpreted the results. M.S. and J.H.C. wrote and revised the manuscript. All authors read and approved the manuscript.

## EXTENDED DATA FIGURE LEGENDS

**Extended Data Fig. 1:**
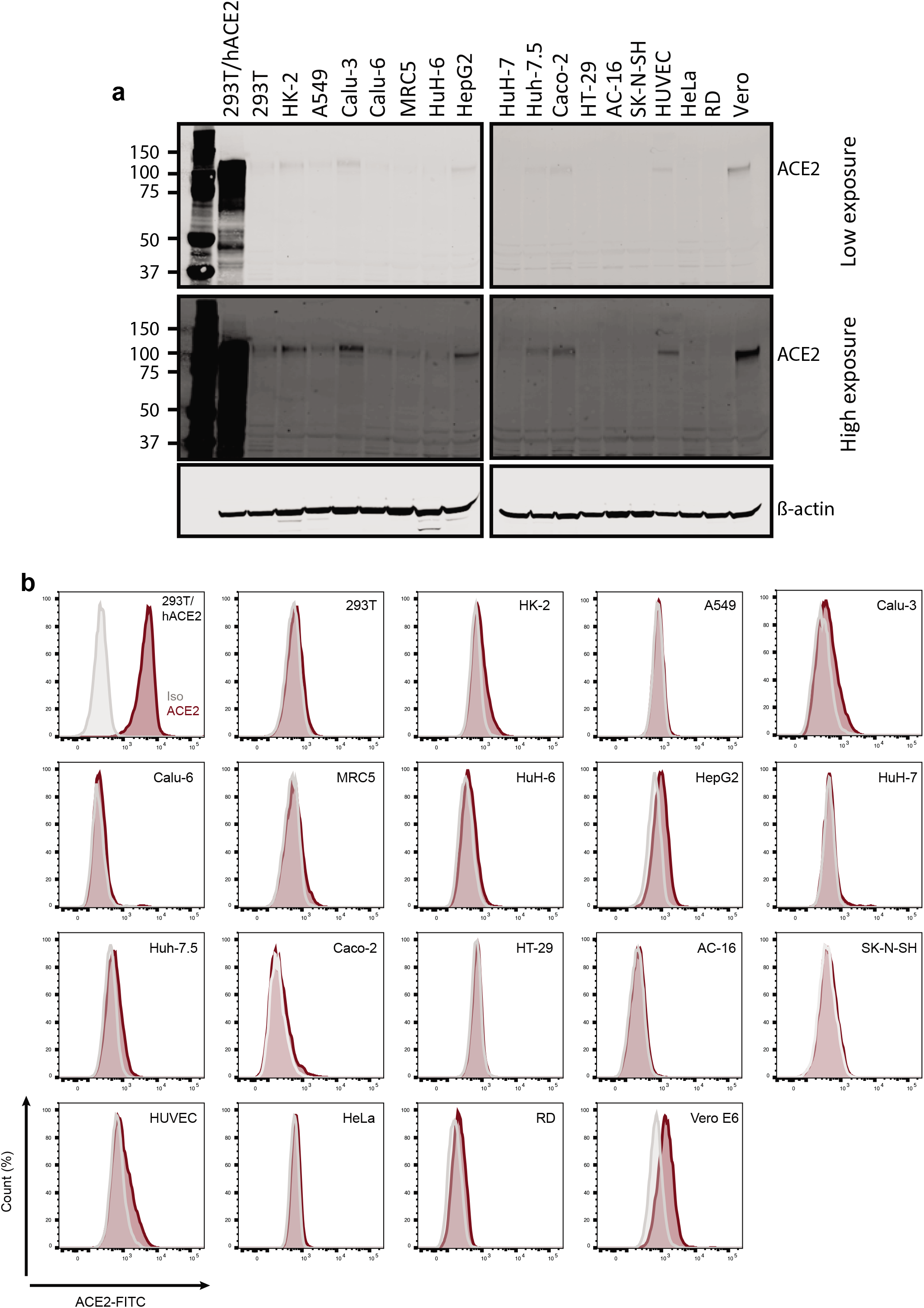
Most human cell lines have low endogenous expression of ACE2. **a**, The ACE2 expression was examined by Western blot. The two different exposure times are shown. β-actin served as a loading control. **b**, Cell surface expression of ACE2 was tested by flow cytometry. Binding of the control IgG is shown in grey histograms while that of the anti-ACE2 antibody in red histograms.

**Extended Data Fig. 2:**
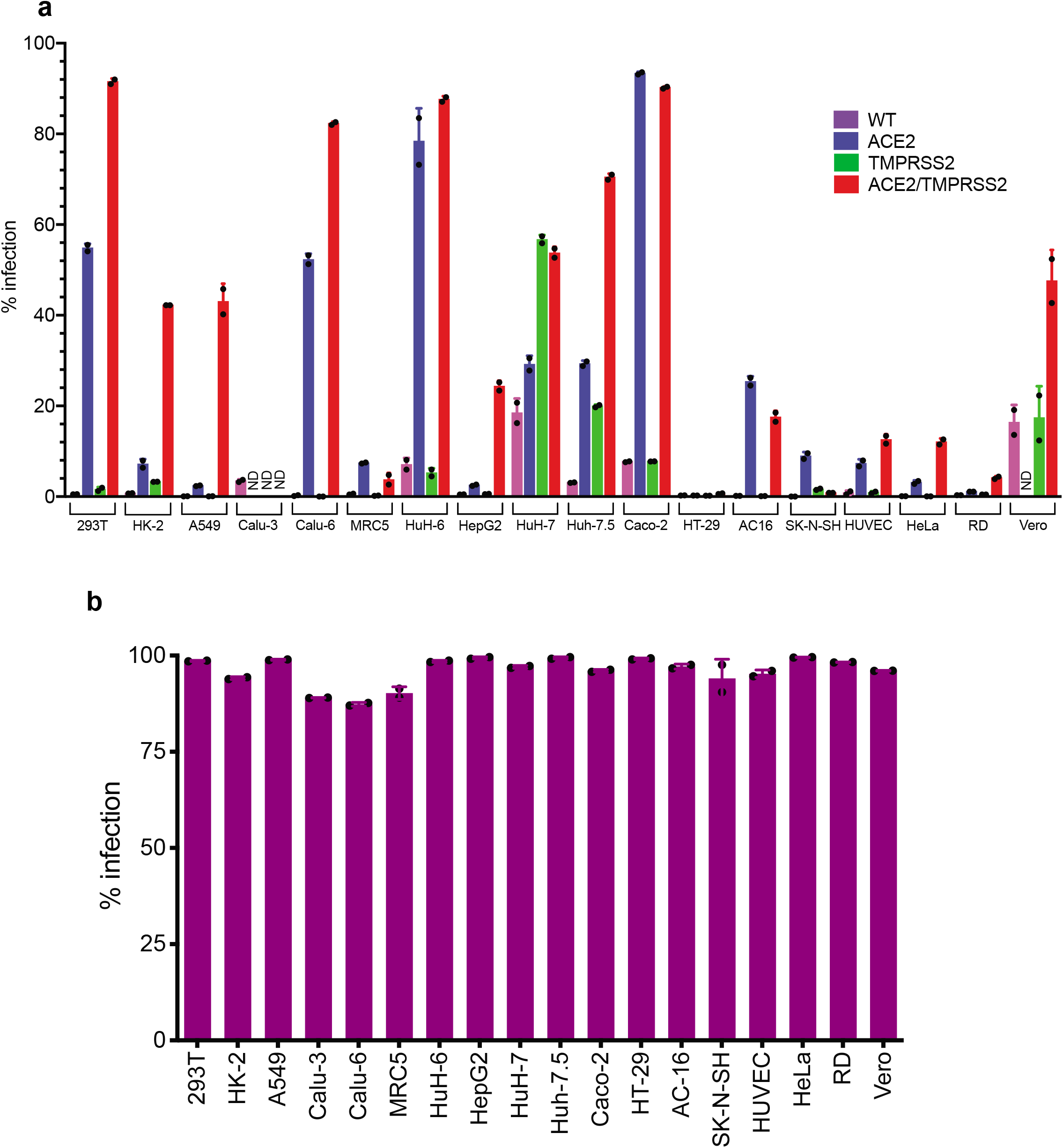
ACE2/TMPRSS2-expression enhances SARS-CoV-2 entry. The indicated cells, engineered to overexpress ACE2 and TMPRSS2, were infected with VSV pseudotyped with either the SARS-CoV-2 spike protein (**a**) or the VSV glycoprotein (**b**). Twenty-four hours later, the infection efficiency was tested by flow cytometry for GFP, encoded by the VSV genome. The mean ± standard deviation of two biological replicates is plotted (n = 2).

**Extended Data Fig. 3:**
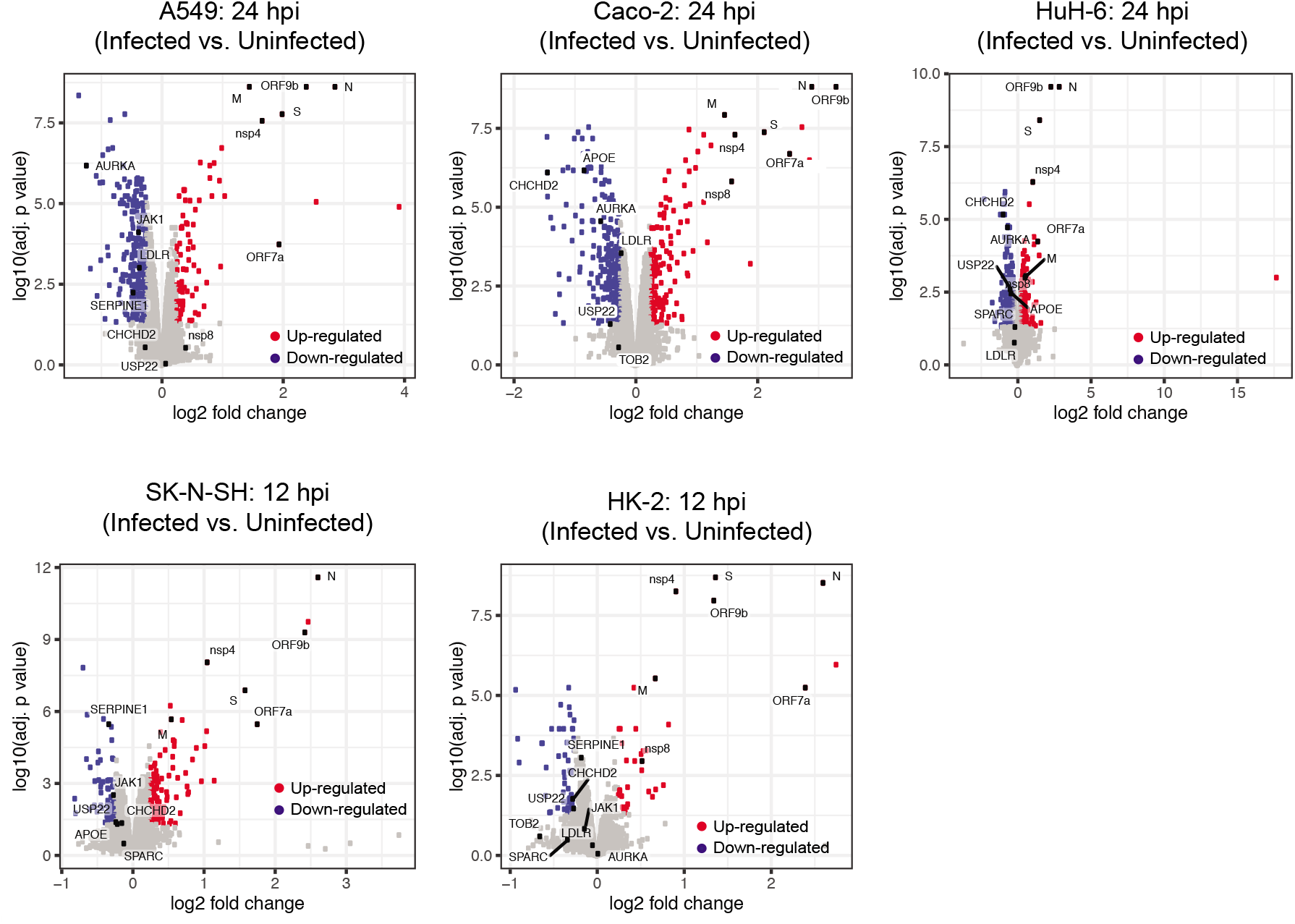
Volcano plots of proteins differentially regulated in different cell lines. The proteins enriched in infected cells are shown in red, while those depleted upon infection in blue color. Black color is used for the proteins labeled with their names.

**Extended Data Fig. 4:**
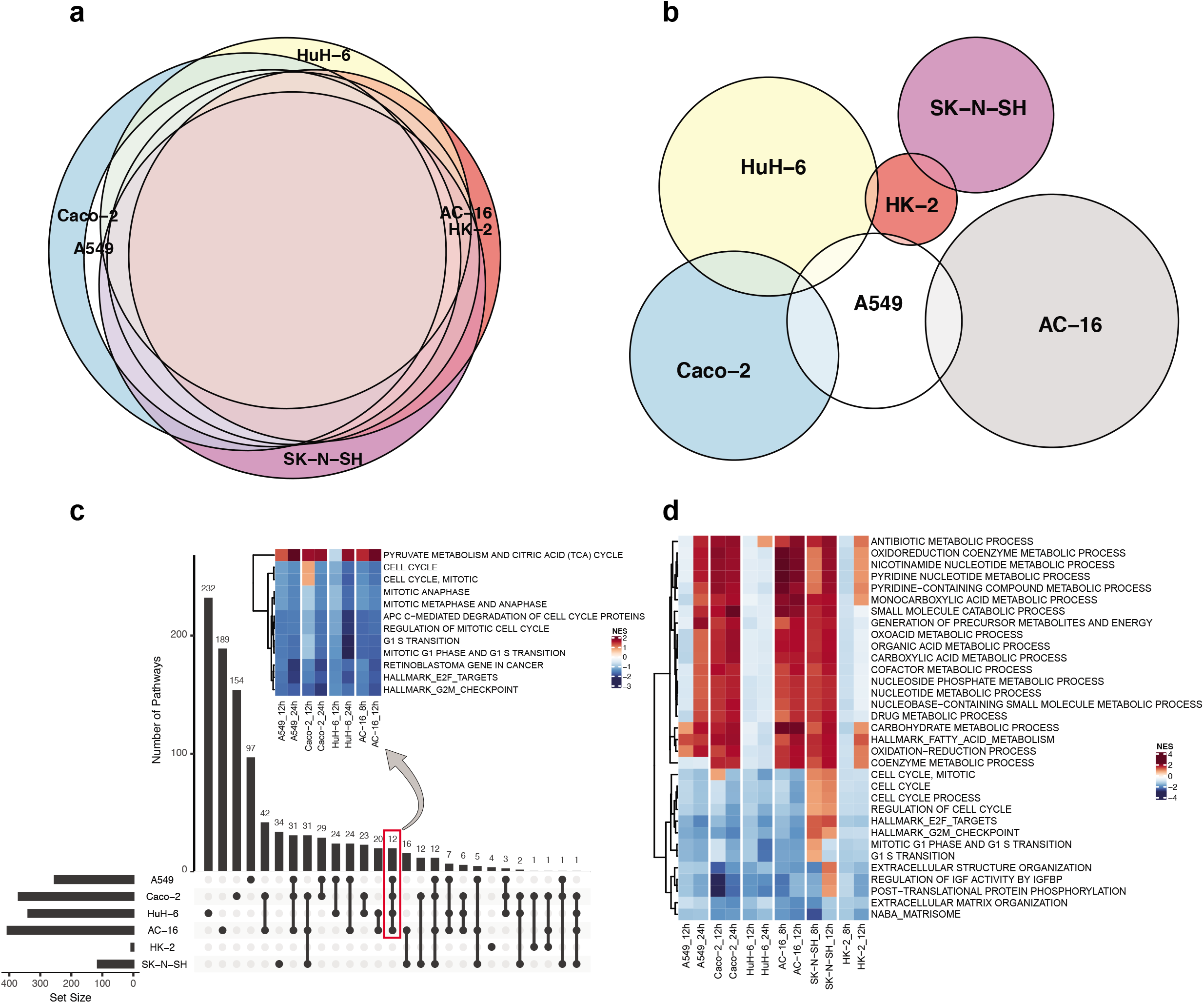
Cellular proteins and pathways commonly targeted by SARS-CoV-2 in multiple cell lines. **a,** Overlap of proteins quantified across six cell lines. **b**, Overlap of proteins regulated across cell lines in a SARS-CoV-2-dependent manner. **c**, Comparative analyses of cellular processes and signaling pathways altered in different cell lines following SARS-CoV-2 infection. Also shown is the heatmap visualization of cellular pathways commonly regulated in four of the six cell lines. Positive and negative enrichment of functional terms is based on normalized enrichment scores (NES). **d**, Heatmap visualization and list of highly differentially regulated pathways and processes in different cell types, identified by Enrichr from the Reactome database (FDR < 0.1).

**Extended Data Fig. 5:**
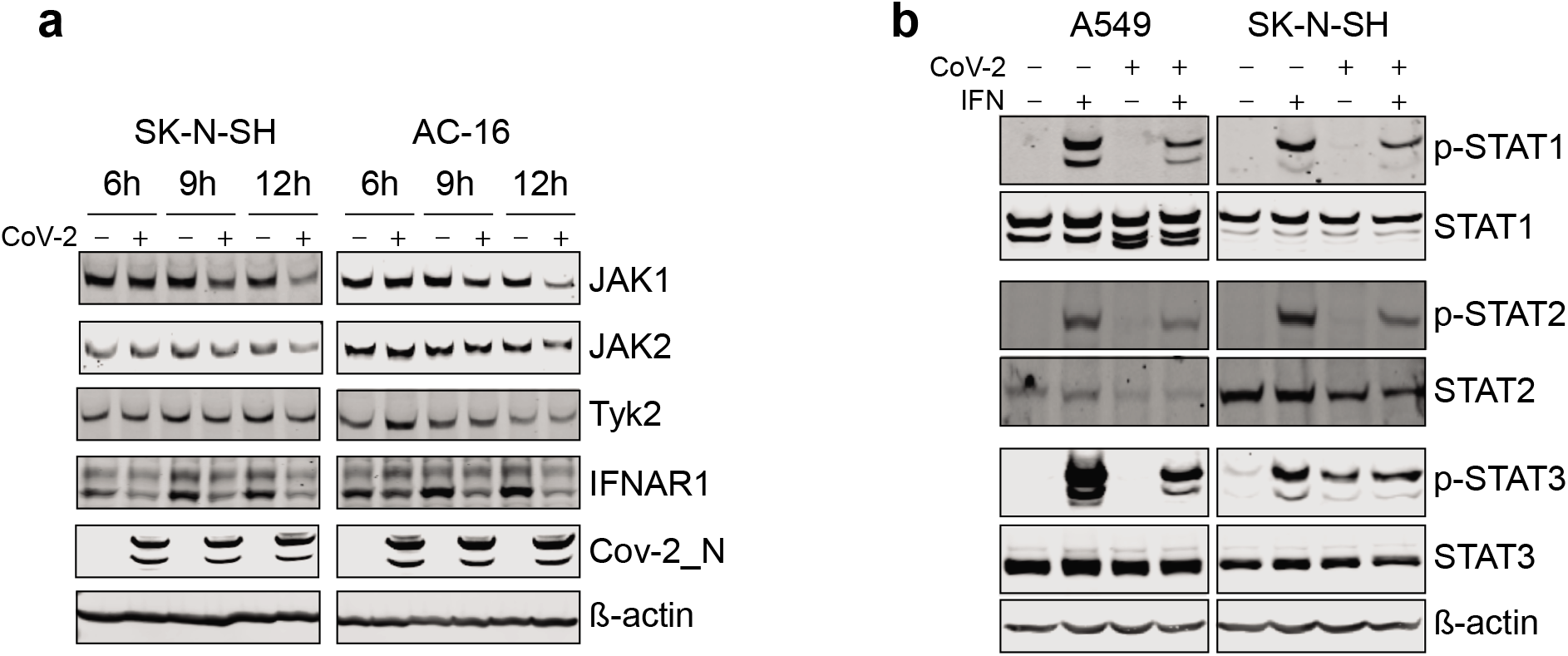
IFN response pathway is impaired in SARS-CoV-2-infected cells. **a,** SK-N-SH and AC-16 cells were infected with SARS-CoV-2 at an MOI of 1 and harvested at 6, 9, and 12 hpi, followed by detection of the indicated proteins by Western blot. **b**, A549 and SK-N-SH cells were infected with SARS-CoV-2 (MOI of 1) for 24h and 12h, respectively, or left uninfected, and then treated with human IFN alpha-2a (1 nM) or, as a negative control, with vehicle (PBS) for 30 min, followed by Western blot for the indicated proteins.

**Extended Data Fig. 6:**
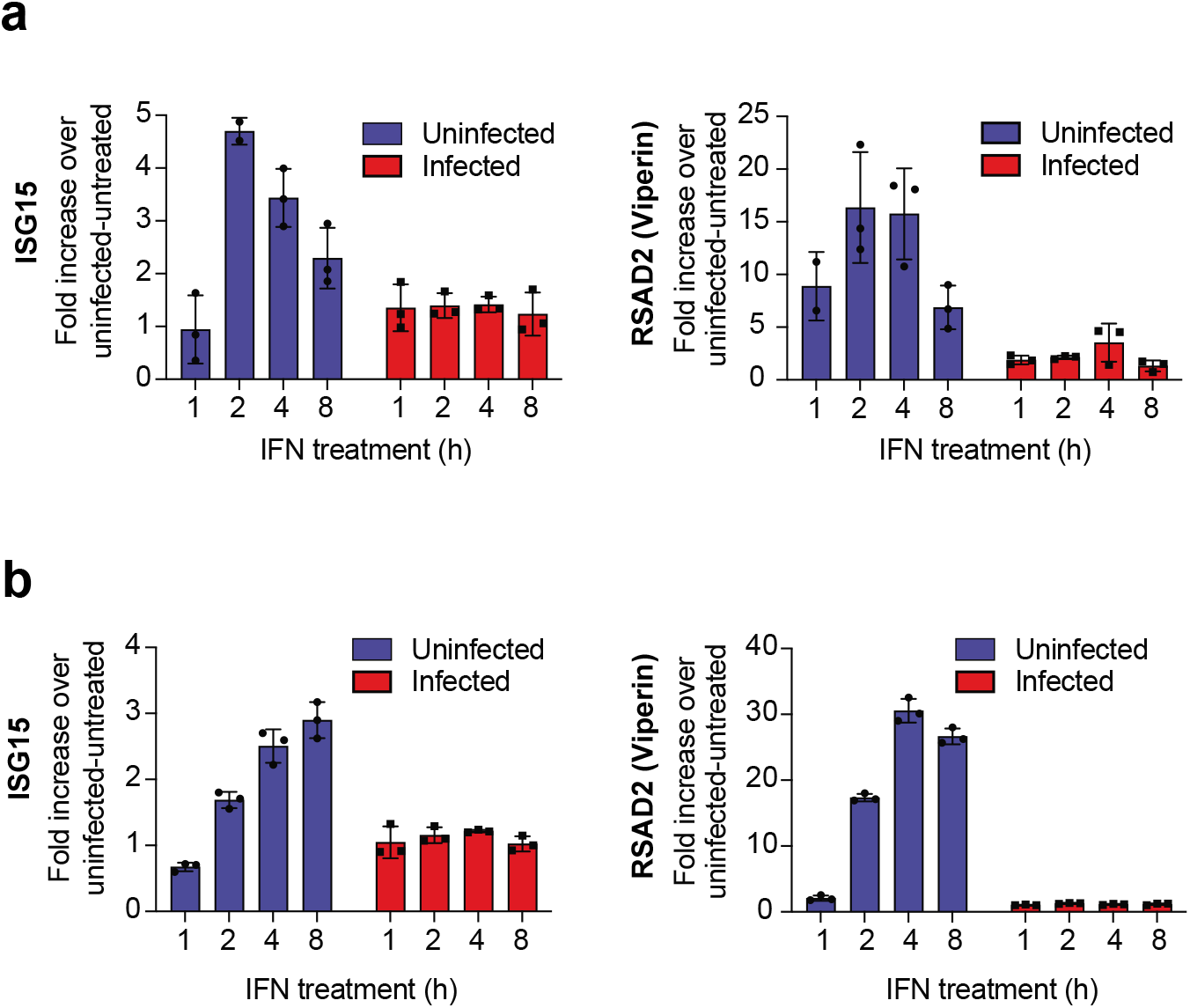
IFN does not cause transcriptional induction of ISGs in SARS-CoV-2-infected cells. **a, b,** A549 (**a**) and Caco-2 (**b**) cells, infected with mock or SARS-CoV-2 for 24h, were treated with IFN (0.1 nM) for 1, 2, 4, or 8h and the RNA levels of the indicated ISGs were measured by RT-qPCR. The values were normalized against RPS11 that served as a housekeeping gene. The data are plotted as mean ± standard deviation of three biological replicates (n = 3).

**Supplementary Excel File: List of differentially regulated proteins in SARS-CoV-2-infected cells.** Time course proteomics data of mock and SARS-CoV-2-infected A549, Caco-2, HuH-6, AC-16, SK-N-SH, and HK-2 cells. The list of proteins was derived from the normalized MaxQuant ProteinGroups.txt files. Shown are the protein groups filtered at a modest cutoff of log_2_ fold change ≥ 0.25, FDR < 0.05.

## Notes

### Competing Interest Statement

The authors have declared no competing interest.

